# Cancer cells resist mechanical destruction in the circulation via RhoA-myosin II axis

**DOI:** 10.1101/601039

**Authors:** Devon L. Moose, Benjamin L. Krog, Lei Zhao, Tae-Hyung Kim, Sophia Williams-Perez, Gretchen Burke, Lillian Rhodes, Marion Vanneste, Patrick Breheny, Mohammed Milhem, Christopher S. Stipp, Amy C. Rowat, Michael D. Henry

## Abstract

During metastasis cancer cells are exposed to potentially destructive hemodynamic forces including fluid shear stress (FSS) while *en route* to distant sites. However, prior work indicates that cancer cells are more resistant to brief pulses of high-level fluid shear stress (FSS) *in vitro* relative to non-transformed epithelial cells. Herein we identify a mechanism of FSS resistance in cancer cells, and extend these findings to mouse models of circulating tumor cells (CTCs). We show that cancer cells acutely isolated from primary tumors are resistant to FSS. Our findings demonstrate that cancer cells activate the RhoA-myosin II axis in response to FSS, which protects them from FSS-induced plasma membrane damage. Moreover, we show that the myosin II activity is protective to CTCs in mouse models. Collectively our data indicate that viable CTCs actively resist destruction by hemodynamic forces and are likely to be more mechanically robust than is commonly thought.

## INTRODUCTION

Circulating tumor cells (CTCs) are a blood-borne intermediate in the metastatic cascade necessary for colonizing distant organ sites. Tumors may generate millions of CTCs per day, but seminal work in cancer biology has established the concept of “metastatic inefficiency” whereby only a small fraction of these CTCs go on to generate clinically observable metastases^1–3^. CTCs exist in a fluid microenvironment quite distinct from that of the solid tumor; in the circulation, these cells are exposed to various biological and mechanical stresses that may lead to their demise, including detachment from extracellular matrix, removal from the trophic factors within the primary tumor, newfound contact with the immune system, and exposure to hemodynamic forces^4,5^. Hemodynamic stresses include fluid shear stress (FSS), shear and compressive stresses due to deformation in the microcirculation and, under some circumstances, traction stresses generated by adherence to the endothelium^6^. Circulatory FSS ranges across 4-orders of magnitude, from less than 1 dyn/cm^2^ in lymphatic vessels and the microcirculation to over 1000 dyn/cm^2^ in turbulent flows in the heart and in certain pathological settings^7–10^. Because cancer cells derived from solid tissues appear to lack adaptations in membrane and cytoskeletal features that allow blood cells to withstand hemodynamic forces^11^, it has often been suggested that CTCs are mechanically fragile relative to blood cells. Indeed, a number of studies indicate that many CTCs are dead or dying^12–15^. However, it is not clear whether death of CTCs is a consequence of the biological and mechanical stresses outlined above or the methods by which CTCs are isolated. It is also possible that many CTCs arrive in the circulation as dead or dying cells, having been passively shed from tumors ^15^. Thus, whether viable CTCs are mechanically fragile is still a matter of speculation.

Other experimental evidence suggests that CTCs may be mechanically robust. For example, studies in mouse models indicate that cancer cells injected into various vascular compartments survive their initial exposure to the circulation, with 85-98% of injected cells viable and arrested in the microcirculation within minutes following injection^2,16–19^. Association of CTCs with blood components such as platelets or CTC clusters may, in principle, afford mechanical protection to CTCs, but there is little direct evidence to support this^20^. Cell intrinsic mechanisms may also contribute to CTC survival in response to mechanical challenges. For example, the mechanosensitive pannexin-1 channel mediates survival in response to cell deformation in the microvasculature^21^. Moreover, we found that, unlike their non-transformed epithelial counterparts, cancer cells are remarkably resistant to brief pulses of high-level FSS^22^, and these findings have since been confirmed and extended by others^23–25^. These studies define that the FSS resistance phenotype is: 1) Evident in cancer cell lines from diverse histologies; 2) Conferred by the presence of transforming oncogenes including *ras, myc* and *PI3K*; 3) Requires the presence of extracellular calcium; and 4) It involves the actin cytoskeleton and the activity of Rho kinase^22,25^. More recently, we found that exposure to brief pulses of FSS results in cell stiffening, suggesting that this stress-stiffening response of cells is related to the FSS resistance phenotype ^26^. A role for cellular mechanical properties in FSS response is further substantiated by reports that lamin A/C contributes to FSS resistance^23^. These data led the hypothesis that cancer cells may actively resist destruction by hemodynamic forces, via changes in cellular mechanics. However, the details of the mechanism underlying FSS resistance have not been described, and whether this feature is functionally relevant for CTCs has not been determined.

## RESULTS

### Viable primary tumor cells are resistant to FSS

Prior studies on FSS resistance analyzed established cancer cell lines, many of which are derived from metastatic tumors^22–25^. To address the possibility that FSS resistance is an artifact of cell culture or the result of metastatic selection, we examined the effects of FSS in tumor cells freshly isolated from a genetically-engineered mouse model of prostate cancer and from human melanoma patient-derived xenografts (PDXs). We exposed the cell suspensions to 10 brief (∼1msec) pulses of FSS (τ_max_=6400 dynes/cm^2^) and assessed FSS resistance after every second pulse, as previously described^22^. Prostate epithelial cells from the tumor-bearing mice displayed greater FSS resistance than those from their wild-type counterparts (Figure 1A). Additionally, we found no difference in FSS resistance between cells isolated from PDX generated from primary and metastatic lesions (Figure 1B). These data demonstrate that FSS resistance is a feature of cancer cells within the primary tumor and not the result of selection during tissue culture or metastasis. Moreover, we observed that FSS resistance was elevated in Pten/Trp53 double knockout vs. Pten only knockout mice (Figure 1A). Given that Pten/Trp53 knockout prostate tumors are known to be markedly more aggressive than their Pten only knockout counterparts^27,28^, the observed difference suggests that FSS resistance may influenced by tumor aggressiveness.

**Figure 1:**
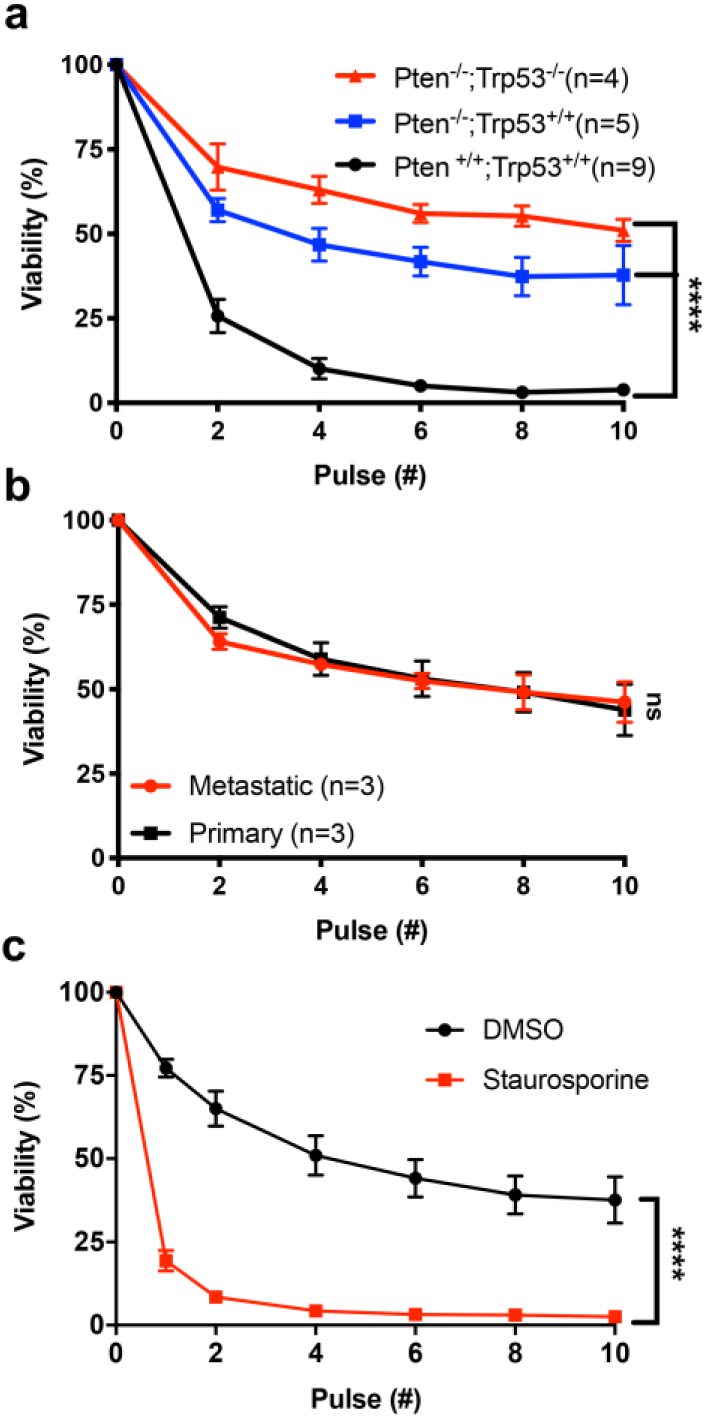
Viable tumor cells are resistant to FSS. A) Effects of 2–10 FSS pulses on the viability of freshly isolated prostate cancer cells from tumor-bearing mice (Pten^-/-^ and Pten^-/-^;Trp53-/-) and on wild-type prostate epithelial cells (****p<0.0001, 2-way ANOVA). B) Effects of 2–10 FSS pulses on cells derived from primary and metastatic human melanoma PDX tumors (p>0.05, 2-way ANOVA). C) Effects of staurosporine treatment vs. vehicle only (DMSO) on FSS resistance (****p<0.0001, n=4; 2-way ANOVA, n=3).

Given that a number of studies have shown that many CTCs are dead or in the process of dying, and our observations on FSS resistance indicate that the latter is an active cellular process, we hypothesized that apparent CTC fragility may be consequence of cell death. To test this, we induced apoptosis in PC-3 prostate cancer cells by treating them with staurosporine (4µM, 4 hr; confirmed by Annexin V staining) and evaluated FSS resistance (Figure S1A). The staurosporine-treated cells were markedly more sensitive to FSS than their untreated counterparts; most were dead after 2 pulses (Figure 1C). We also used ionomycin to induce necrotic-like cell death; we found that after a single pulse of FSS nearly all of the ionomycin treated cells had been destroyed (Figure S1A&B). These data reaffirm that FSS resistance is an active cellular process.

### Cancer cells are intrinsically resistant to small (<12nm) membrane disruptions induced by FSS

Our previous study had demonstrated that exposure to FSS results in membrane damage, detected as uptake of the membrane impermeant dye propidium iodide (PI), in otherwise viable PC-3 prostate cancer cells^22^. To characterize the plasma membrane damage produced by our FSS protocol further, we exposed PC-3 prostate cancer cells to FSS in the presence of fluorescent dextrans of increasing molecular weight and evaluated cellular uptake and viability by flow cytometry. As the size of the dextran probe increased from 3,000 to 70,000 MW, progressively less was taken up by viable cells (Figure 2A). After 10 pulses of FSS, less than 5% of cells showed evidence of uptake of 70,000 MW dextran, whereas ∼30% had taken up 3,000 MW dextran, consistent with the fraction of viable cells that took up PI (668 MW) (Figure 2B). Since the Stokes diameter of 70,000-molecular weight dextran is ∼12nm^29^, exposure to FSS in our protocol resulted in relatively small, and reparable holes in the plasma membrane of cancer cells. Thus, the FSS resistance of cancer cells could reflect a greater capacity to repair the plasma membrane in response to damage or, alternatively, a greater intrinsic resistance to membrane damage than that of non-transformed cells.

**Figure 2:**
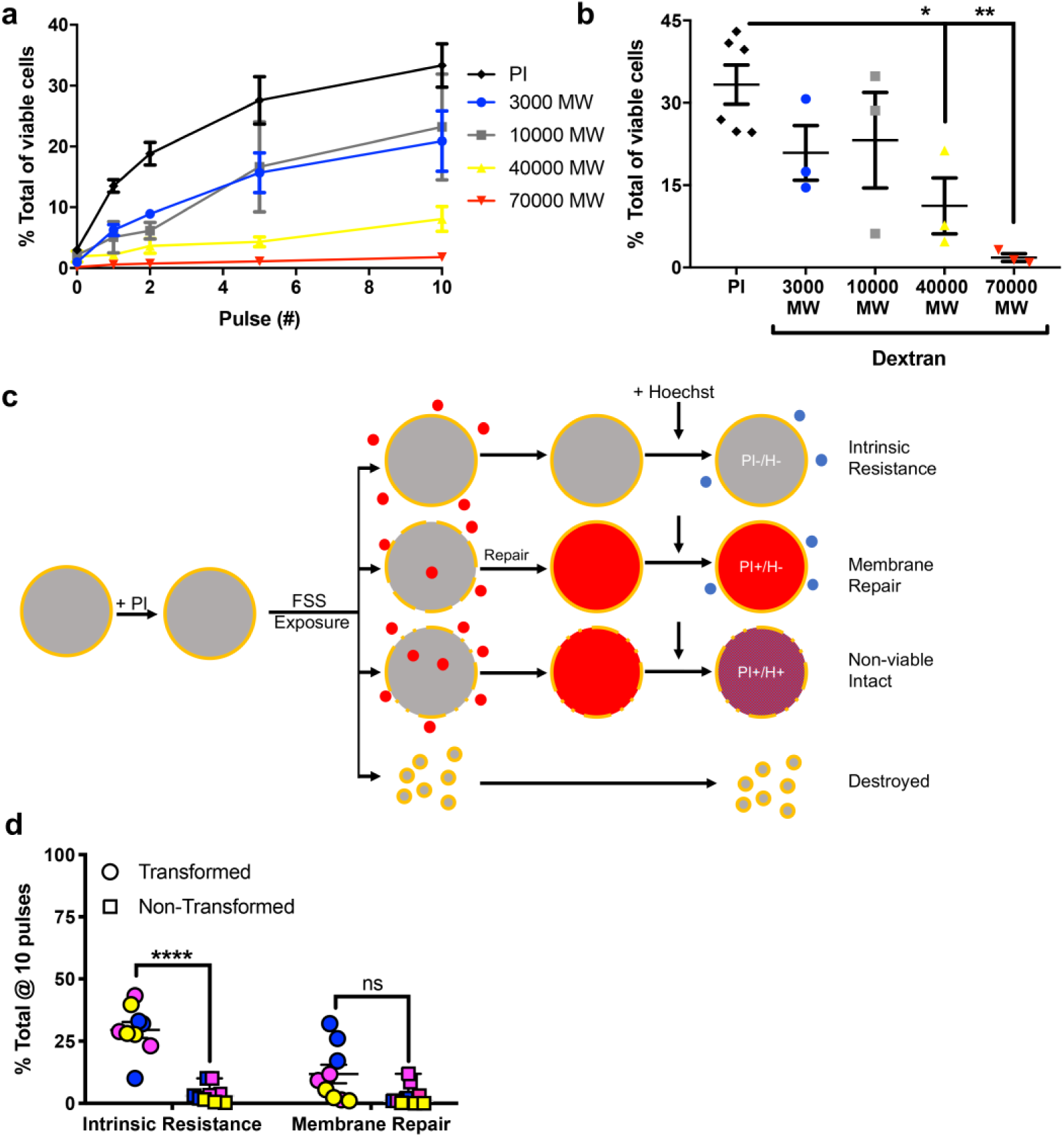
Cancer cells resist FSS-induced nanometer-scale disruption of the plasma membrane. A) Uptake of dextrans of various size and of propidium iodide (PI) in PC-3 cells exposed to FSS. B) Uptake after 10 pulses of FSS. Uptake of both 40,000 and 70,000 MW dextrans decreased in comparison to that of PI (*p<0.05 and **p<0.01, respectively; 1-way ANOVA with Bonferroni correction), C) Schematic of flow cytometry assay for evaluation of intrinsic resistance and membrane repair after exposure to FSS. Shown is sequence of exposure to PI, FSS, and Hoechst dye, making it possible to identify cells that have intrinsic resistance to membrane damage (PI^-^Hoechst^-^) and cells that undergo membrane repair (PI^+^Hoechst^-^). D) Effects of 10 FSS pulses on intrinsic resistance and membrane repair in transformed cancer cells (PC-3, blue circle; MDA-MB-231, pink circle; TCCSUP, yellow circle) and non-transformed cells (PrEC-LH, blue square; MCF-10A, pink square; Primary urothelial, yellow square) (***p<0.001, t-test, n=3/cell line). Membrane repair after 10 pulses varied among cell lines and did not differ significantly by transformation status (p>0.05, t-test, n=3/cell line).

To distinguish between the roles of repair and intrinsic resistance, we developed a flow cytometry-based assay that measures the fraction of viable cells in which membrane damage is repaired or in which no membrane damage occurs (Figures 2C and S2A). Before exposing the cells to FSS, we briefly added PI, and afterwards we added counting beads and the viability dye Hoechst. Because membrane damage must be repaired rapidly in order to maintain cell viability, cells that repair damage to the plasma membrane will be PI-positive but Hoechst-negative. Intrinsically resistant cells, i.e., those that do not experience membrane damage, will be negative for both dyes. Utilizing this assay, we tested for the capacity of three pairs of cancer and non-transformed cells to resist and repair FSS-induced plasma membrane damage. In the case of the transformed cell lines, PC-3, MDA-MB-231, and TCCSUP, all exhibit a consistent level of intrinsic resistance (33.4%, 31.8%, and 31.8%, respectively) after 10 pulses of FSS (Figure 2D; S2B, D, F, H). This level of intrinsic resistance was substantially elevated over that in non-transformed cells, with all three such cell lines, PrEC LH, MCF-10A, as well as primary urothelial cells, having less than 5% after 10 pulses of FSS (Figure 2D; S2C, E, G, H). Of the transformed cells that were evaluated, only PC-3 cells had a significant repair fraction (33.5%). Both MDA-MB-231 and TCCSUP displayed only minimal evidence of repair (7.5% and 3.0%) (Figure 2D, S2I). Non-transformed MCF-10A cells showed an appreciable repair fraction during initial pulses that was diminished after 10 pulses, indicating that their susceptibility to damage outweighs their ability to repair damage. Both PrEC-LH and primary urothelial cells exhibited only marginal membrane repair (Figure 2D; S2I). These data demonstrate that, although membrane repair may contribute to FSS resistance of some transformed cell lines, cancer cells are consistently distinguished from non-transformed cells by intrinsic resistance to membrane damage.

### Resistance to FSS depends on RhoA-myosinII axis

To determine the cause of increased intrinsic resistance to FSS-induced damage in cancer cells, we considered previously published findings suggesting that cell stiffness may contribute to FSS resistance. Those studies demonstrated that: 1) PC-3 cells stiffen in response to FSS^26^, and 2) utilization of F-actin and Rho kinase inhibitors can sensitize cancer cells to FSS^22,25^. Based on these findings, we hypothesized that members of the Rho-family GTPases, which regulate actin dynamics and actomyosin contractility, are activated in response to FSS. To test this hypothesis, we utilized RhoA/C and RAC1 GTP pulldown assays. We found that in PC-3 cells both RhoA and RhoC were activated in response to FSS, while active RAC1 levels were similar(Figure 3A-B; S3A-D). We then knocked down RhoA and RhoC in a PC-3 derivative cell line and found that only RhoA knockdown cells had decreased FSS resistance (Figure S3E). Utilizing the RhoA activation assay, we found that FSS exposure activates RhoA in both MDA-MB-231 and TCCSUP cells. This demonstrates that cancer cells from diverse tissues respond to FSS in a similar manner by activating RhoA (Figure 3A-B). Since RhoA activation typically leads to increased levels of F-actin^30,31^ we assayed for cortical F-actin levels in PC-3 after two pulses of FSS exposure by using imaging flow cytometry. This revealed a robust increase in the formation of cortical F-actin, which is consistent with increased RhoA activity (Figure 3C; S4A-B).

**Figure 3:**
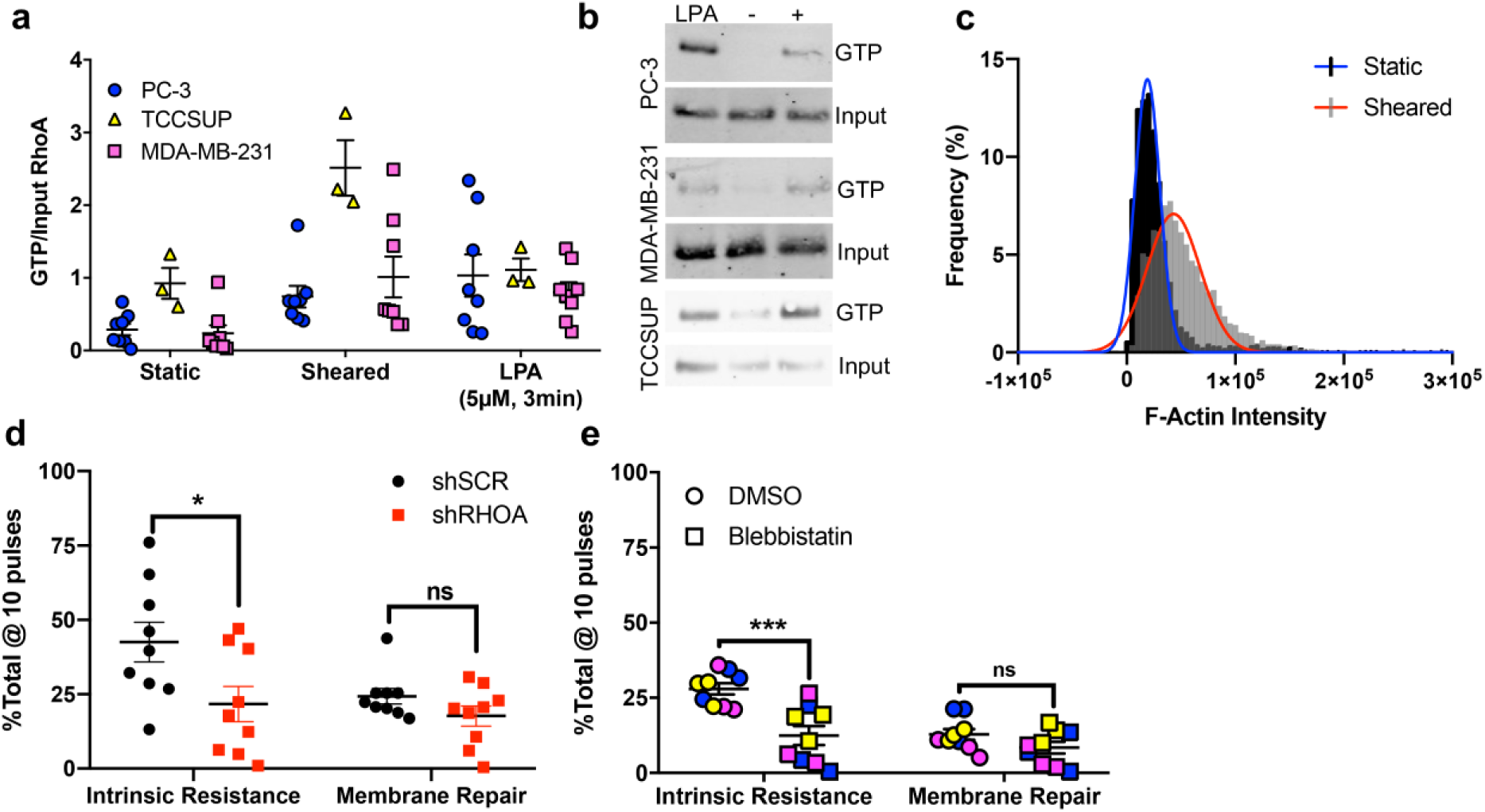
The RhoA-myosin II axis is required for intrinsic resistance to FSS. A) Effects of FSS pulses on cortical F-actin levels in PC-3 cells (****p<0.0001, Welsh’s t-test). B) RhoA activation in PC-3 (*p<0.05, n=8, Welch’s t-test), MDA-MB-231 (*p<0.05, n=8, Welch’s t-test), and TCCSUP (*p<0.05, n=3, Welch’s t-test) cells in suspension under static conditions (static), after 2 pulses of FSS (sheared), and in response to lysophosphatidic acid (LPA; positive control). C) Representative western blots of active RhoA from pull-down assay. D) Effects of 10 FSS pulses on intrinsic resistance (*p<0.05, t-test, n=9) and membrane repair in cells treated with control and RhoA-targeting shRNAs (p>0.05, n=9 t-test). E) Effects of blebbistatin on intrinsic resistance (***p<0.01, t-test, n=3/cell line) and membrane repair (p>0.05, t-test, n=3/cell line) in response to FSS exposure.

To determine whether the RhoA-myosin II axis enables cancer cells to maintain elevated intrinsic FSS resistance, we knocked down RHOA in PC-3 cells using RNA interference and the flow cytometry assay described above (Figure 2C). We found that the intrinsic FSS resistance was lower in RHOA knockdown vs. scramble control cells (Figure 3D; S5A-C). We then used blebbistatin to inhibit myosin II activity in PC-3, MDA-MB-231 and TCCSUP cells and found that the fraction of intrinsically resistant cells was significantly reduced (Figure 3E; S5D-H). After 10 pulses of FSS exposure, the membrane repair fraction was not significantly affected by either genetic perturbation of RHOA or pharmacologic (blebbistatin) inhibition of myosin II (Figure 3D-E, S5A-E). However, blebbistatin treatment did alter the membrane repair during the course of FSS exposure (Figure S5E). We also utilized inhibitors that target Rho kinase and myosin light-chain kinase, and found that these too sensitized PC-3 cells to FSS (Figure S5F,G). Importantly, at the exposures used in these studies, none of these pharmacologic agents influenced the viability of cells prior to FSS exposure (Figure S5H). This indicates that the effects of these compounds on FSS resistance are due to specific inhibition of this mechanism rather than a secondary consequence of cytotoxicity. Taken together, these studies establish the importance of the RhoA-myosin II axis in intrinsic FSS resistance.

### Inhibition of myosin II sensitizes cancer cells to hemodynamic forces

We next sought to determine whether the results above implicating myosin II activity in FSS resistance are relevant to CTCs *in vivo*. To this end, we adapted a mouse model that had been developed to assess the trapping of viable cells in the lung microvasculature following their injection via the tail vein ^32^. PC-3 cells were labeled with distinct viable fluorescent dyes, pretreated with either blebbistatin or DMSO (vehicle control), and mixed with 15μm fluorescent microspheres prior to injection. Following injection, the mice were euthanized within 3 min to assess the short-term effects of the circulation on trapping of intact cells in the lung microvasculature (Figure 4A). We found that the number of intact PC-3 cells lodged in the lungs was significantly lower in the blebbistatin treated vs. DMSO group (49.71±1.57% vs. 59.01±2.67%) (Figure 4B-E; Table S1). Within this experiment we also swapped the dyes used for each treatment condition and found that the results obtained were independent of the dye label used (Figure 4F). This indicated that it was the blebbistatin treatment that led to the reduction in the number of intact cancer cells lodged in the lung microcirculation. We reasoned that this effect could have been due to either destruction of blebbistatin-treated cells *en route* to the lung or an increase in their capacity to traverse the lung microvasculature. To discriminate between these possibilities, we took advantage of the expression of firefly luciferase in the cells tested; we found that the activity of cell-free luciferase correlated with the reduction in viability of the cells exposed to FSS *in vitro* (Figure S6A-C). By measuring the cell-free luciferase detected in the plasma of injected mice, we were able to directly quantify the number of cells destroyed during this brief exposure to the circulation. Destruction was 2-fold higher for the blebbistatin-treated samples vs. their DMSO-treated counterparts (6.2±0.6% vs. 3.8±0 .6%) (Figure 4G; S6D). Although these differences in cell lodgment and destruction were statistically significant, we note that the magnitude of the change was moderate; this could be a consequence of the very brief (few minutes) period of time during which the cancer cells were exposed to the circulation.

**Figure 4:**
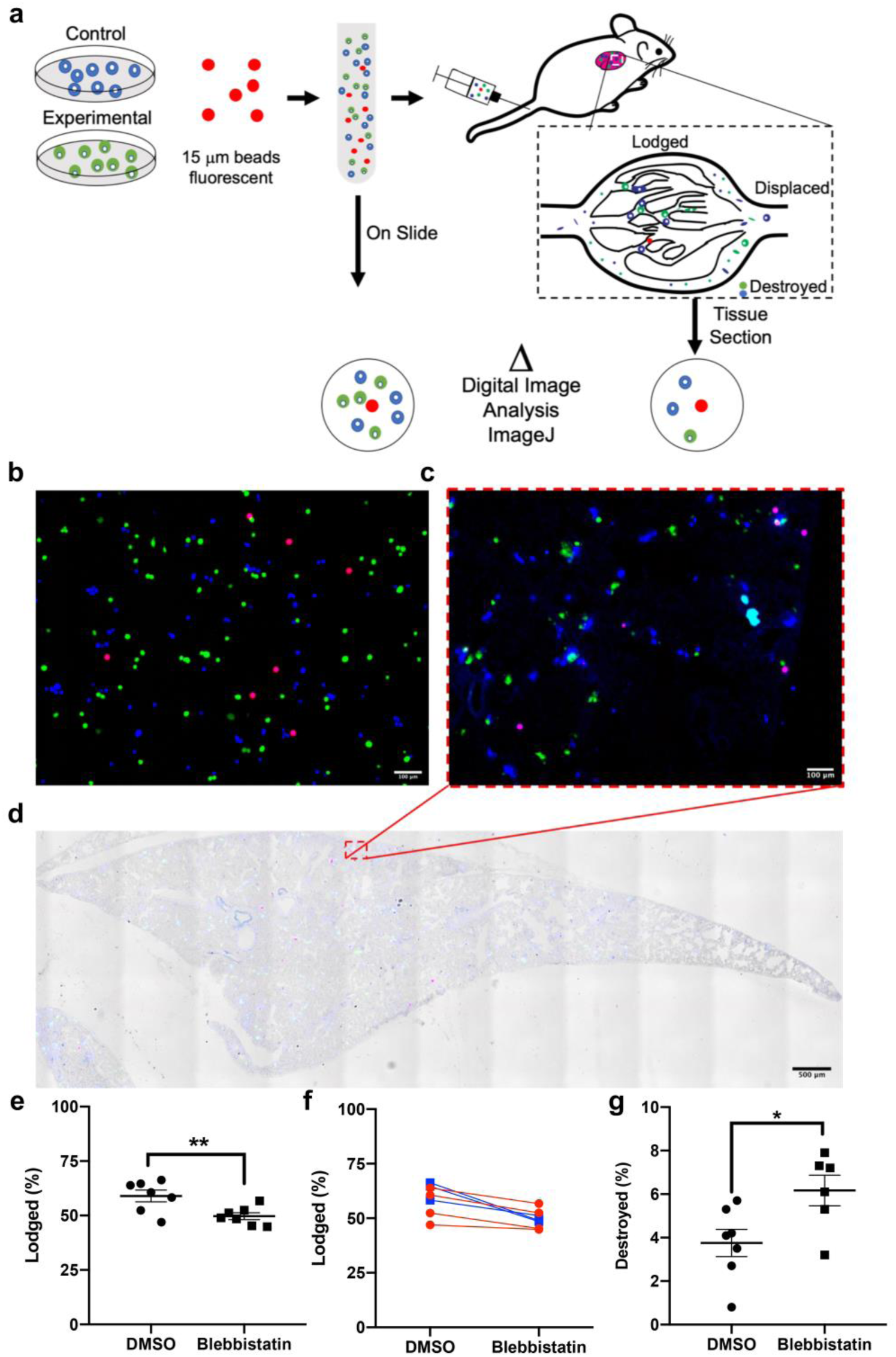
Myosin II activity is required for the lodgement of intact CTCs in the lung and reduces cellular destruction immediately following entry into the circulation. A) Schematic of the experiment, with DMSO control cells (shown in blue) and blebbistatin-treated cells (shown in green) mixed with microspheres labeled red prior to injection into mouse for assessment of lodgement in the lung. B-D) Cells treated with blebbistatin are pseduocolored green and cells treated with DMSO pseduocolored blue; beads are red. B) Representative image of uninjected control sample (“On Slide”). Bar, 100µm C-D) Lung sections with cell types and microspheres labeled as depicted in A. C) Bar, 100µm; D) Bar, 500µm E-G). Quantitation of effects of blebbistatin treatment on E-F) lodgement of intact cells (**p<0.01, n=7, paired t-test) and G) cell destruction (*p<0.05, n=6,7, Welch’s t-test). F) Graph of the lodgement data for each mouse, with the lines colored to group each dye condition (p>0.05, 2-Way ANOVA, n=7).

To test whether inhibition of myosin II can affect the survival of CTCs over a longer period of time in a mouse model of metastasis, we employed an orthotopic model of metastatic prostate cancer that we had previously developed ^33^. Utilizing this model, we treated tumor bearing mice with a single dose of blebbistatin (2.5 mg/kg) and measured CTC number in cardiac blood 3 hours later (Figure 5A). Neither the overall tumor burden nor tumor growth differed between the two groups (Figure S7A-B). We observed a dramatic reduction in CTC number in mice treated with blebbistatin vs. the vehicle control (Figure 5B; S7C). Importantly, over a range of exposures bracketing the *in vivo* exposure level, blebbistatin was not directly cytotoxic to the prostate cancer cells, indicating that the effects of blebbistatin on prostate cancer CTCs cannot be accounted for by direct toxicity (Figure S7D). Moreover, treatment of non-tumor bearing mice with blebbistatin did not affect the number of CD45^+^ cells detected in blood (Figure 5C), indicating that blebbistatin was not toxic to leukocytes. This suggests that the RhoA-myosin II axis promotes FSS resistance specifically in cancer cells rather than in blood cells more generally. Taken together, our data strongly indicate that targeting of the RhoA-myosin II axis could sensitize CTCs to destruction by hemodynamic forces.

**Figure 5:**
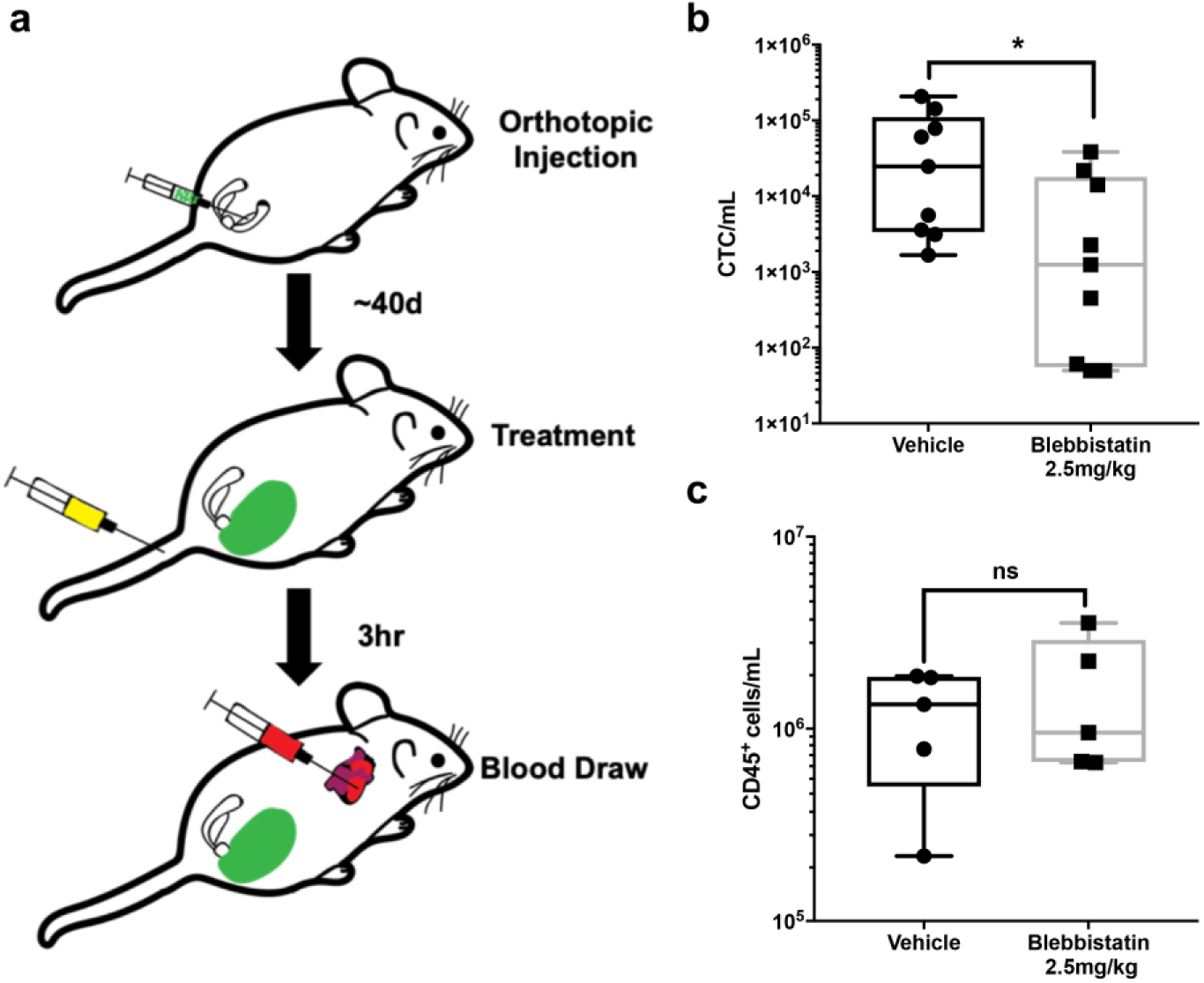
Myosin II activity is required to maintain steady-state levels of CTCs. A) Schematic of the experiment, with orthotopic tumor cell injection, blebbistatin treatment at ∼40 d, and blood draw to test CTC number 3 hr later. B) Effects of blebbistatin treatment on CTC number (∼1200 vs ∼25000 CTC/mL) (*p<0.05, n=9/group, Mann-Whitney U test). C) Effects of blebbistatin treatment on CD45^+^ cells (1.3×10^6^ vs 9.5×10^5^) (p>0.05, n=5/group, Mann-Whitney U test).

## DISCUSSION

Cancer cells rely on the circulatory system to spread beyond the primary tumor and regional lymph nodes. This journey presents a mechanical challenge to those originating in solid organs, in part because FSS in the circulation is orders of magnitude higher than that experienced due to interstitial flow in tissues^34^. Here we show that cancer cells actively resist FSS-induced damage to the plasma membrane via a mechanism involving the RhoA-myosin II axis. Our study challenges the long-held notion that CTCs are inherently fragile. Although, numerous studies have shown that an appreciable fraction of CTCs are dead or in the process of dying CTCs following their isolation^12–15^, the cause of death of these cells remains unclear, as they are subjected to both biological and mechanical insults^4,5^. Our data indicate that dead and dying cancer cells are highly susceptible to destruction by FSS, suggesting that biological fragility precedes mechanical fragility. Thus, we conclude that mechanical destruction in the circulation is does not contribute significantly to metastatic inefficiency, supporting a number of previous studies^2,16–19^.

Precisely how CTCs experience FSS and other hemodynamic forces in the circulation is not clear^34^. The best available evidence indicates that the periods during which CTCs circulate freely last only a few seconds, and that they are interspersed between much longer periods during which they are trapped in the microcirculation due to size restriction. However, CTCs vary in size, and some may be less prone to lodgment in the microcirculation^35^. It is possible that RhoA-myosin II activation protects CTCs from damage by hemodynamic forces as shown here, but that this activity is modulated dynamically as CTCs traverse the microcirculation, with cells that are less stiff more readily “crawling” through narrow channels^36^. In our studies, the reduced lodgment of blebbistatin-treated cells in the lung microcirculation was not evenly balanced by the observed increase in destruction, suggesting that some of these more compliant blebbistatin-treated cells may have avoided lodgment in the microcirculation. Passage through the heart may expose CTCs to brief, high-magnitude FSS, most similar to that used our *in vitro* model. Using this model we showed that intrinsic resistance to plasma membrane damage, as opposed to elevated membrane repair, most consistently distinguished cancer cell lines (bladder, breast and prostate) from their non-transformed epithelial counterparts. In PC-3 prostate cancer cells, we observed that exposure to FSS produced small (<12nm) holes in the plasma membrane. These small holes may be rapidly repaired via tensional forces in the plasma membrane^37^.

We show that FSS resistance is a phenotype in cancer cells isolated directly from primary tumors. It is not an artifact of cell culture, nor the product of metastatic selection. Our previous study indicated that FSS resistance is conferred by various transforming oncogenes, such as *ras, PI3K* and *myc*. Although it remains to be determined how these oncogenes accomplish this, plausible mechanisms linking *ras* and *PI3K* with RhoA activity have been reported^38,39^. Our finding that exposure to FSS activates RhoA indicates that the development of resistance to FSS is an adaptive response. Previously, we found that exposure to FSS results in cell stiffening ^26^, which likely reflects activation of the RhoA-myosin II axis. We do not yet know exactly what triggers RhoA activation in response to the mechanical stimulus of FSS. Our demonstration that extracellular calcium is essential for FSS resistance in cancer cells^22^ suggests that calcium influx, through either small membrane disruptions that we detect or possibly a mechanosensitive ion channel, activates RhoA or triggers myosin II contractility directly. Although we find that the RhoA-myosin II axis is important for FSS resistance, our efforts to genetically or pharmacologically inhibit this pathway did not completely ablate FSS resistance. The fact that withdrawal of extracellular calcium achieves this end suggests that other, as yet unidentified, calcium-dependent processes also influence FSS resistance. Moreover, cell stiffness is a complex phenotype involving not only cytoskeletal contractility but also the properties of the glycocalyx, composition of the plasma membrane, osmotic state and nuclear structure and stiffness, with the latter having already been implicated in FSS resistance^23,40^. Thus other mechanisms are likely to modulate FSS resistance in cancer cells.

Finally, our demonstration that a blockade of myosin II activity leads to an increase in the destruction of CTCs has interesting implications. It is notable that a reduction of over an order-of-magnitude was evident within 3 hours of treatment with blebbistatin, and that the number of CD45^+^ leukocytes was not affected. These outcomes suggest that the myosin II-mediated promotion of FSS resistance is specific for cancer cells. It is not yet clear if FSS resistance represents a new anti-cancer target. Intervening in the steps of metastatic dissemination for therapeutic benefit is regarded as challenging as it may occur before cancer diagnosis^41^. However, metastatic seeding is likely an ongoing process in cancer patients^42^ and during surgical and diagnostic procedures, CTCs may be liberated and could be targeted at that time^43^. Should the reduction in CTCs be found to translate to reduced metastasis, it is possible that the targeting of mechanisms that lead to FSS resistance will be effective in anti-metastatic therapy. Although blebbistatin is not a drug candidate in humans, other agents such as Fasudil, a ROCK inhibitor, may be effective in this regard^25^. Moreover, because the activation of RhoA-myosin II has been implicated in other aspects of cellular behavior that could promote metastatic colonization, e.g., extravasation, cell invasion and cell survival^44^, the anti-metastatic benefits of inhibiting this pathway may be pleiotropic. This also raises the interesting possibility that exposure to FSS ”primes” cells for subsequent metastatic behavior.

## Material and Methods

### Cell lines

All cell lines were cultured according to protocols provided by the suppliers. Cancer cell lines were obtained from ATCC. Primary bladder epithelial cells were acquired from Lonza. The non-transformed prostate epithelial cell line PrEC LH was acquired from Dr. William Hahn (Dana Farber Cancer Institute). The GS689.Li cell line was derived from the human prostate cancer cell line PC-3 that were twice selected *in vivo* for increased metastatic potential ^45^. The identity of this cell line as a derivative of PC-3 cells was validated by short tandem repeat (STR) analysis (IDEXX). Cells of both the PC-3 and GS689.Li lines were cultured in DMEM/F12 (Gibco) supplemented with 10% FBS (Atlanta Biologicals, S11150) and 1X NEAA (Invitrogen). The breast cancer line MDA-MB-231, 293FT, and human epithelial kidney line HEK293T were cultured in DMEM with 10% FBS, and 1X NEAA. PrEC-LH cells were cultured in prostate epithelial cell growth medium (Lonza, CC-3166), primary bladder epithelial cells were cultured in prostate epithelial basal medium (ATCC, PCS-440-030) using the corneal epithelial growth kit (ATCC, PCS-700-040), and cells of the non-tumorigenc breast cancer cell line MCF-10A were cultured in DMEM/F12 with 5% horse serum (Invitrogen, 16050), 20 ng/mL EGF (Sino Biological, 10605HNAE250), 0.5 μg/mL hydrocortisone (ACROS, 352450010), 10 μg/mL insulin (Sigma, 91077C), and 1x Pen/Strep (Invitrogen, 15070). PC-3, GS689.Li, and MDA-MB-231 cells were transfected with an integrating retrovirus that encodes the firefly luciferase gene using procedures previously described ^46^. RhoA knockdown lentivirus and its control were generated by transfecting 293FT cells with VSVG and the PLKO.1 vector, with the latter of containing a short hairpin RNA (shRNA) targeting either RhoA or a non-target sequence (Sigma), using the Polyfect reagent (Qiagen). The cells were selected for antibiotic resistance 2 days after transduction, and knockdown was evaluated by western blot analysis.

### Chemical Reagents

Cells were treated with blebbistatin (20μM, 3 hr) (Selleckchem, S7099), fasudil (5μM, 24 hr) (Selleckchem, S1573), ML7 (20μM, 1 hr) (Sigma, I2764), ionomycin (10μM, 30 min) (Sigma, I9657), or staurosporine (4μM, 4 hr) (Selleckchem, S1421) prior to exposure to fluid shear stress (FSS). For mouse studies, blebbistatin (-) (Selleckchem, S7099) was injected once intravenously, at a dose of 2.5 mg/kg.

### Fluid shear stress assay

Cells were suspended in DMEM without FBS at a concentration of 5×10^5^cells/mL and exposed to FSS at a flow rate of 250 μL/s in a 30 Ga ½ ” needle using a 5-mL syringe and a PHD1000 syringe pump (703006). For cells expressing luciferase, viability was determined by bioluminescence imaging ^22^. For cells that did not express luciferase, the CellTiter-Blue reagent (Promega) was used. In all cases, viability was determined by normalizing signal to the static control (pulse 0).

### Flow cytometry

Data were collected using a Becton Dickinson LSR II flow cytometer and analyzed using the FLOWJO platform (FLOWJO, LLC).

#### Evaluation of Intrinsic Resistance and Membrane Repair

To detect damage to the plasma membrane, 1.75 µg/mL propidium iodide (PI) was added to a cell suspension before it was subjected to FSS. The size of plasma membrane disruptions was measured by adding 10 µM of 3, 10, 40, or 70 kDa FITC-dextran (LifeTechnologies; D1821, D3305, D1844, D1823) to the suspensions prior to FSS exposure. Samples were collected at “pulse 0” (static control), as well as immediately after pulses 1, 2, 5, and 10. They were washed with FACS buffer (1x PBS, 0.5% BSA, 0.1% NaN_3_), centrifuged at 500g for 3 min, then resuspended in the FACS buffer with 6 µg/mL Hoechst 33258 (Thermo Fisher; H3569) and either 2-µm YG microspheres (Polysciences; 17155-2) or 15-µm scarlet microspheres (Invitrogen; F8843). For calculating the percentage of cells in each fate, the ratio of intact cells to the counting beads was first calculated. For each pulse evaluated, the intact ratio was normalized to the static control sample (pulse 0). The normalized intact ratio was than multiplied by the percentage of cells for each fate (PI^-^Hoechst^-^, PI^-^Hoechst^+^, PI^+^Hoechst^+^, and PI^+^Hoechst^-^).

#### Staining for Annexin V

PC-3 cells were treated with DMSO or 4 µM staurosporine for 4 hr. Prior to FSS exposure (pulse 0), 500 µL samples were taken and assessed for Annexin V expression. The samples were washed with staining buffer (BioLegend, 420201), centrifuged at 500 g for 3 min, and resuspended in 100 µL of Annexin V Binding Buffer (BioLegend, 422201) with allophycocyanin (APC)-conjugated Annexin V (1:20; BioLegend, 640920) for 30 min. 400 µL of Annexin V binding buffer with Hoechst 33258 and 15-µm scarlet microspheres was added prior to analysis.

#### Staining for CD45

Non-tumor bearing male mice were treated with 2.5 mg/kg of either blebbistatin or vehicle. At 3 hr after treatment, blood was collected by terminal cardiac draw, and 500 µL of blood was mixed with 10 mL of red blood cell (RBC) lysis buffer (150mM NH_4_Cl, 10mM NaHCO_3_, 1.3 mM EDTA), centrifuged at 500 g for 3 min, and resuspended in FACS buffer. Samples were blocked in FACS for 30 min and incubated with anti-mouse CD45 (1:100, BioLegend 103110) for 1 hr. Samples were washed 3 times with FACS buffer and suspended to the volume initially collected, in FACS buffer containing a specific concentration 15-µm scarlet microspheres.

#### Analysis of circulating tumor cells (CTCs)

Blood samples collected for flow cytometry analysis were processed using RBC lysis buffer, centrifuged at 500 g for 3 min, then suspended to the volume initially collected, in FACS buffer containing either 2-µm YG microspheres (Polysciences; 17155-2) or a known concentration 15-µm scarlet microspheres. The initial concentration of the 2-µm YG microspheres was calculated using the known concentration of 15-µm scarlet microspheres. Hoechst 33258 was used to distinguish between live and dead circulating EGFP^+^ tumor cells. The volume analyzed was determined by dividing the number of microspheres analyzed by their concentration (microspheres/mL) in the sample. The cell concentration (cells/mL) was calculated from the cell count and the volume analyzed. ^47^

### Rho GTPase activity assay

For both the static and FSS-exposed samples, cells were released from the tissue-culture dish with 0.25% trypsin (Gibco 25200-056), suspended in media containing FBS, centrifuged at 300 g for 5 min, resuspended in 5 mL of DMEM, and held in suspension for 30 min prior to exposure to two pulses of FSS at 250 μL/sec. Lysophosphatidic acid (LPA) positive controls were serum starved overnight in DMEM with 5 mg/mL bovine serum albumin (BSA), then treated with 5 μM LPA (Sigma, L7260) for 3 min prior to lysis. To detect GTP bound RhoA or RhoC, cells were lysed using Rho activity buffer (50mM Tris, 500mM NaCl, 50mM MgCl2, 1% Triton X-100, 0.1% SDS) and the volume was normalized such that 800-1000 μg of protein was used for each pulldown. To detect GTP bound RAC1, cells were lysed using RAC activity buffer (50mM Tris, 500mM NaCl, 50mM MgCl2, 1% Triton X-100) and 800-1000 μg of protein used for each pulldown. For each pulldown sample, 5% was saved for use as the input control and the rest was incubated with recombinant protein (Rho binding domain of Rhotekin for RhoA/C or of PAK1 for RAC1) (> 30 μg) for 45 min at 4 °C. Samples were washed 3 times with 1 mL of RhoA activity buffer and centrifuged at 4 °C and 500 g for 3 minutes.

### Western blotting

Samples were loaded onto SDS-polyacrylamide gels (NuPage 4-12% Bis-Tris Protein Gels, Novex) and transferred to PVDF membranes (Immobilon-FL), which were incubated in primary antibody (1:500 mouse anti-RhoA, ARH04, Cytoskeletal Inc.; 1:1000 mouse anti-β-tubulin, Developmental Studies Hybridoma Bank; 1:1000 mouse anti-Rac1 BD Bioscience; or 1:1000 rabbit anti-RhoC, D40E4, Cell Signaling) overnight. The blots were subsequently incubated with secondary antibody (1:10000 goat-anti-mouse; Rockland, 610-731-124; 1:10000 goat anti-mouse Jackson Labs, 715-036-151; or 1:20000 goat anti-rabbit LiCOR, 925-68071) and signal was assessed using either an Odyssey (LI-COR) or a ChemiDocX (BioRad) system.

### Quantification of Cortical F-actin

To measure the density and organization of F-actin after shear stress, we exposed PC3 cells to 2 pulses of FSS. To identify dead and damaged cells, we added propidium iodide (1.75 µg/ml) to the cell suspension before FSS. Immediately following FSS, we fixed cells with 1% paraformaldehyde for 20 min. After washing with PBS, cells were stained with 1:500 phalloidin-AlexaFlour488 (ThermoFisher, A12379) and 1:500 DRAQ5 (ThermoFisher, 62251) in staining buffer (2% FBS in PBS, v/v) for 30 min. Cells were then washed with PBS and resuspended with 80 µL of staining buffer for imaging flow cytometry (ImageStream®X Mark II, EMD Millipore) at the UCLA Flow Cytometry Core Facility of Eli and Edythe Broad Center of Regenerative Medicine and Stem Cell Research. To quantify cortical F-actin levels, we used the ImageStream Data Analysis and Exploration Software (IDEAS) (Amnis Corporation). The cortical region was defined by subtracting the Erode mask (8 pixel) from the Dilate mask (1 pixel) of the brightfield channel.

### Animal models

All procedures were approved by the University of Iowa Animal Care and Use Committee (protocol #’s 1302028, 4121221, 5121574).

#### Genetically engineered mouse model of prostate cancer

The C57BL/6J-Try^c-2J^/J ROSA26-LSL-Luc;Pbsn-cre;Pten^fl/fl^ mice were generated as described previously ^27^. The C57BL/6J-Try^c-2J^/J ROSA26-LSL-Luc; Pbsn-cre; Pten^fl/fl^; Trp53^fl/fl^ mice were generated by breeding Pten knockout mice with B6.129P2-Trp53^tm1Brn^/J counterparts (Jackson Laboratory) and backcrossing progeny for 6 generations onto the C57BL/6J-Tyr^c-2J^ background ^48^. When mice of the test generation were 25 weeks of age, prostate tissue was collected from those with Cre-activated luciferase that were homozygous for either the wild-type allele, floxed Pten, or floxed Pten and floxed Trp53, at 25 weeks of age.

#### Patient-derived xenograft (PDX) model

Patient tumor samples to be used in PDX models were collected and processed by the University of Iowa College of Medicine Tissue Procurement Core under IRB# 200804792. Tissue collection and distribution was performed in accordance with the guidelines of the University of Iowa Institutional Review Board. Briefly, following collection tumor samples were placed in DMEM containing 5% FBS, 1X Pen/Strep, and 1% fungizone. For initiation of PDX tumors, patient tumor samples were processed by mincing until they passed through an 18 G needle easily. The cells were then suspended in a 1:1 mixture of DMEM (Gibco) and Matrigel (Corning, 354230), and subcutaneously injected into 2 or 3 NOD.Cg-Prkdc^scid^ Il2^rgtmWjl^/SzJ (NSG) (Jackson Laboratory) mice. For the propagation of samples, tumors were dissociated into single-cell suspensions and each mouse was injected subcutaneously with 1×10^6^ cells. Single-cell suspensions were generated by placement of the tumor into a sterile petri dish containing 10 ml collagenase IV solution (9 ml HBSS (Ca^++^- and Mg^++^-free), 1 ml collagenase IV (Worthington Biochemical Corp.), and 5 mM CaCl_2_). The tumor samples were first minced and incubated for 1 hr at 37 °C. Following this, samples were further digested in 10 ml 0.05% trypsin (at 37°C for 10 min), and passed through an 18 G needle 30 times, after which the suspension was filtered using a 70μm cell strainer.

#### Tail-vein metastatic colonization model

Cells were labeled with 1 µM Cell-Tracker Green (Invitrogen; C7025) or Cell-Tracker Red (Invitrogen; C34552) and 2µg/mL Hoechst 33342 (Invitrogen; H1399) for 30 min prior to being mixed with 15-µm microspheres (Invitrogen; F8843) and suspended in PBS. A sample of the mixture was taken for the control to calculate the cell-to-microspheres ratio, after which 200 μL were injected into the tail vein of a Balb/c mouse (Charles River). Mice were anesthetized with isoflurane immediately after injection and terminal cardiac blood collection was performed. Blood was placed in heparinized collection tubes (BD Microtainer, 365974) for further processing. The lungs were inflated by tracheal injection with a 1:1 mixture of OCT and PBS, mounted in OCT, and flash frozen. Lungs were then sectioned at 20-µm thickness and sampled at 200 µm intervals. Whole-lung sections were imaged. Cells were counted and the number of microspheres was calculated by image analysis as described below.

#### Orthotopic prostate xenograft model

5×10^4^ GS689.Li cells were orthotopically injected into the anterior prostate lobe of 8-13 week-old NSG mice as described previously ^49^. As described above, this cell line was derived from the PC-3 parent line by selection for cells that are highly metastatic *in vivo* ^45^. These cells were engineered to express luciferase and enhanced green fluorescent protein (EGFP), which allows for monitoring tumor burden and the detection of CTCs, respectively. Mice were randomized into two groups with similar tumor burden and injected intravenously with 50 µL of either 2.5 mg/kg (-)-blebbistatin or vehicle (15% DMSO, 85% PEG-400). Blood was collected 3 hr after injection by terminal cardiac puncture, and samples were analyzed by flow cytometry as described below.

### Image analysis

Microscopy images were acquired using an Olympus BX61 or a Leica SP8 microscope. For imaging of the lung, a region of interest (ROI) encompassing the entire lung section was selected, and images were collected through 4 fluorescence channels and stitched together using microscope software (LAS X, Leica or cellSens, Olympus). Image analysis was performed using the FIJI software ^50^. The threshold for the detection of fluorescent cells was set as signal >4 standard deviations above the mean fluorescence intensity across the lung section. ROIs were selected using the analyze particle tool in FIJI. The presence of a nucleus in an area positive for cytoplasmic fluorescence was determined by measuring nuclear signal in the ROIs for the fluorescence channels corresponding to Cell-Tracker Red and Green; ROIs were considered to contain a nucleus only if the average signal was greater than the mean + 1 standard deviation of the fluorescent signal for the lung section in the nuclear (Hoescht 33342) channel. The percentage of intact cells lodged in the microvasculature was determined based on the ratio of cells to microspheres in the lungs at p2 vs p0 (pre-injection control sample).

### Bioluminescence imaging

After orthotopic tumors were implanted, their growth was monitored to assess tumor burden by weekly bioluminescence imaging. Imaging entailed intraperitoneal injection of 150 mg/kg of D-luciferin (GoldBio, LUCK), followed 5 min later by the detection of bioluminescence using an AMI X imager (Spectral Instruments Imaging); exposure time was 5 minutes. The AMIView Imaging Software was used to select an ROI including the whole body for quantification of signal intensity.

### Cell-free luciferase assay

#### Assay validation

To validate that cell-free luciferase measurements correlate with the destruction of cells due to FSS, we compared the calculated percentage of cells destroyed by exposure to FSS with the loss of viability as measured with bioluminescence signal (Fig. S9). Specifically, luciferase expressing PC-3 cells were suspended in 90% DMEM and 10% ddH_2_O and then subjected to FSS. To generate a standard curve, we lysed PC-3 cells at a concentration of 5×10^6^ cells/mL in 1% Tween-20 in ddH2O and subsequent dilution the lysate 1:10 with DMEM, and measured the cell-free luciferase signal that correspond to 0, 25, 50, 75, and 100% of cells destroyed. We took 1 mL samples prior to and after 10 pulses of FSS exposure, centrifuged at 1,500 × g for 5 min, and the supernatants were transferred into new tubes. Subsequently, 100μL aliquots were pipetted into wells of a 96-well black-bottom plate (Corning 3915) and 100μL of assay buffer (200mM Tris-HCl pH=7.8, 10mM MgCl2, 0.5mM CoA, and 0.3mM ATP, 0.3 mg/mL luciferin) was placed in each well to measure luciferase activity. The fractional cell viability as measured in luciferase assay of cells exposed to FSS was plotted against a standard curve of the cell-free luciferase signal generated from cell-lysate from a known quantity of cells.

#### Application of cell-free luciferase assay to mouse plasma

A standard curve was generated from data for 5×10^5^ cells lysed with 200 μL 1% Tween-20 in ddH2O. The standard curve was generated using samples in which lysates were titrated into 500 μL of freshly isolated mouse blood, corresponding to the destruction of 2.08×10^5^, 1.04 ×10^5^, and 5.20 ×10^4^ cells. Assuming that BALB/c mice have ∼1.5 mL of blood ^51^, these samples represent ∼33% of the blood pool into which the cells were injected; this number was the basis of normalization of the luciferase signal obtained from the standard curve to the total blood pool. Both the blood for the standard curve and samples collected from mice injected with cells via the tail vein were centrifuged at 2,500 × g for 5 min. Subsequently, 100μL of plasma was collected and combined with 100μL of luciferase assay buffer (200mM Tris-HCl pH=7.8, 10mM MgCl2, 0.5mM CoA, and 0.3mM ATP, 0.3 mg/mL luciferin) for measurement of the luciferase activity. Only blood collected from mice that had fully patent injections were used for analysis. The number of cells destroyed following tail-vein injection was assessed by simple linear analysis using the standard curve.

### Statistical analysis

Data were analyzed using the statistical tests indicated in each figure legend; when appropriate, Bonferroni’s correction for multiple comparisons was used. For each experimental group, the FSS assay was repeated at least 3 times over independent passages. For the F-actin data in fig. 2, the signal intensity was transformed using log_10_. The concentrations of CTCs and CD45^+^ cells in Fig. 4 are presented as medians and interquartile range, the flow analyses for intrinsic resistance and membrane repair in figs. S3, S6, and S7 are presented as stacked bar charts of the component means, and all other data are presented as the mean with SEM indicated by error bars. Statistical significance was set at P<0.05 and all statistical tests were two-sided.

## Acknowledgments

We would like to thank Justin Fishbaugh and Dr. Chantal Allamargot for technical assistance with flow cytometry and microscopy, respectively, and Drs. Christine Blaumueller, Kevin Campbell, Frank Solomon and Dan Welch for helpful comments on the manuscript. Human tumor specimens were obtained through the University of Iowa Melanoma, Skin & Ocular Repository (MaST), an Institutional Review Board-approved biospecimen repository.

### Funding

This project was supported by NIH grants R21 CA179981 and R21 CA196202 to MDH and the University of California Cancer Research Coordinating Committee (CRR-18-52690) to ACR. DLM was supported by Pharmacological Sciences Training grant (2T32-GM0677954-14). Core facilities at the University of Iowa were supported by P30 CA086862 to the Holden Comprehensive Cancer Center.

### Author contributions

Conceptualization: DLM, BLK, MDH; Analysis: DLM, BLK, LZ, THK, PB, ACR, MDH; Investigation: DLM, BLK, LZ, THK, SWP GB, LR; Methodology: DLM, BLK, THK, CS, ACR, MDH; Project Administration: MDH; Resources: MV, MM, CSS; Writing-Original Draft: DLM & MDH Writing-Review & Editing: All authors.

### Competing interests

MDH is a co-founder of SynderBio, Inc.

