## Supplemental Figures for "Cancer cells resist mechanical destruction in the circulation via RhoA-myosin II axis"

**List of supplementary material:**

*Figure S1:* *Dead cancer cells are rapidly fragmented by FSS*
*Figure S2:* *Transformed cells have a higher intrinsic resistance to FSS than their non-transformed counterparts*
*Figure S3* *Exposure to FSS activates Rho and not Rac family GTPases*
*Figure S4* *Exposure to FSS results in increased cortifal F-actin*
*Figure S5* *RhoA-myosin II axis confers elevated intrinsic resistance to FSS*
*Figure S6* *Validation of assay for measuring cell destruction based on levels of cell-free luciferase*
*Figure S7* *Blebbistatin reduces steady-state CTC number*
*Figure S8* *Western blot images*
*Table S1* *Fewer blebbistatin-treated cells lodge in the lung microvasculature*

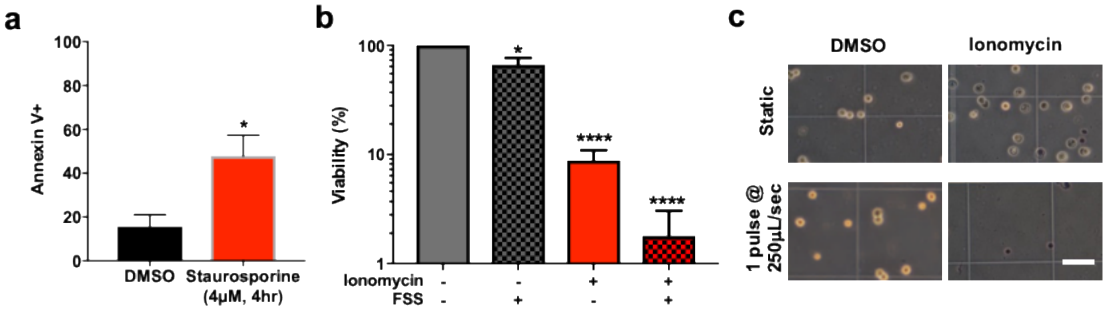


**Figure S1: Dead cancer cells are rapidly fragmented by FSS.** A) Apoptosis in PC-3 cells treated with staurosporine (4μM, 4hr), as assessed by Annenxin V staining (*$p<0.05$, n=4; t-test). B) Effects of ionomycin treatment (10 μM, 30 min) on susceptibility of PC-3 cells to 1 pulse of FSS at 250 μL/s. DMSO is vehicle control for ionomycin treatment. Ionomycin treatment significantly reduced viability (****$p<0.0001$, n=3; 1-way ANOVA with Bonferroni) and sensitized cells to FSS (****$p<0.0001$, n=3; 1-way ANOVA with Bonferroni) while the reduction in viability due to shear stress alone was not as great (*$p<0.05$, 1-way ANOVA with Bonferroni). C) Phase-contrast images of cells before and after ionomycin treatment and/or FSS exposure. Bar, 100mm.

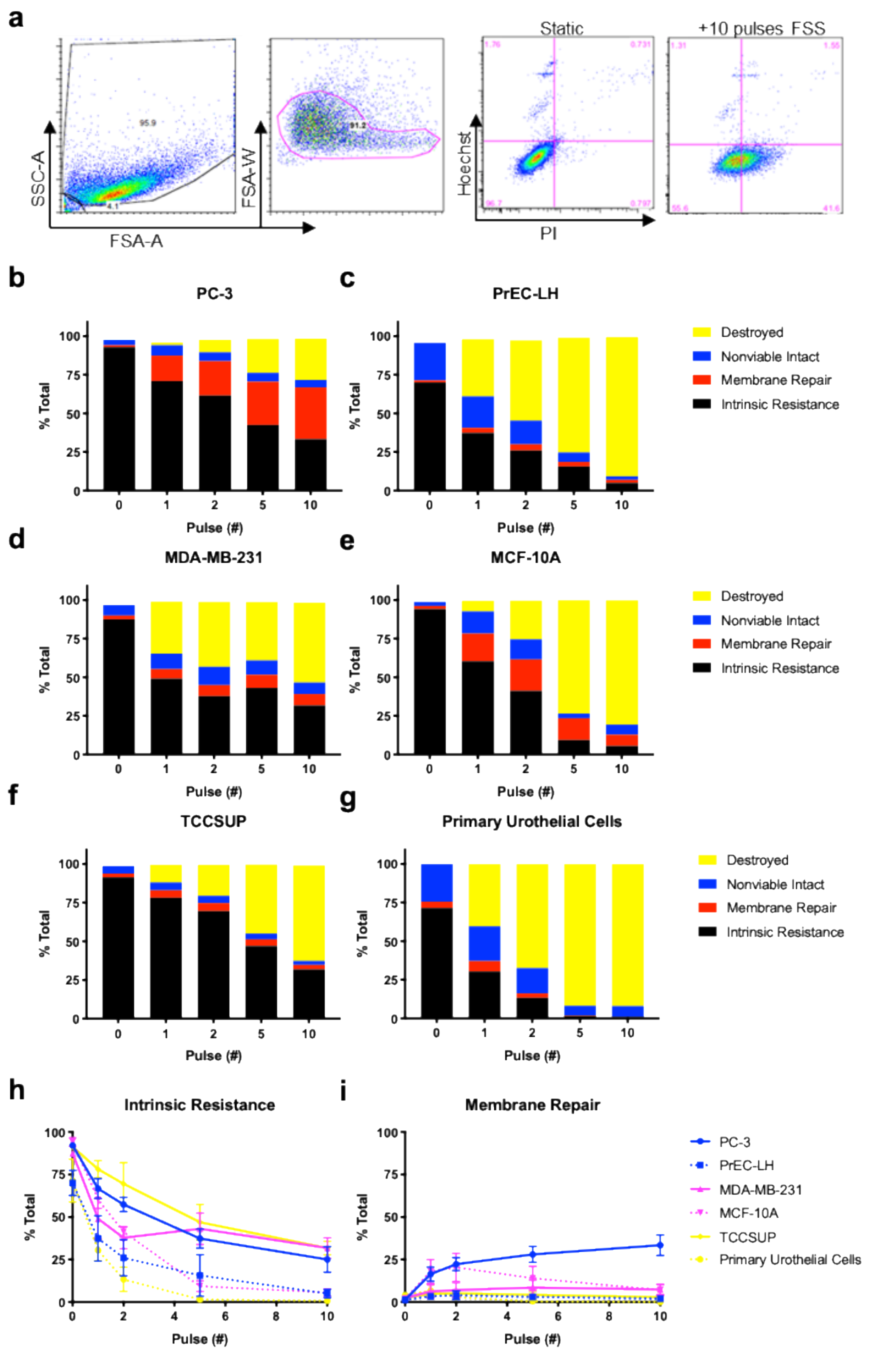


**Figure S2: Transformed cells have a higher intrinsic resistance to FSS than their non-transformed counterparts**. A) Representative gating profile with PI and Hoechst stain for static and sheared PC-3 cells (10 pulses). B-G) Histograms showing population means over the course of exposure for intrinsic resistance, membrane repair, nonviable intact, and destroyed for paired transformed (PC-3, MDA-MB-231, TCCSUP) and non-transformed (PrEC-LH, MCF10A, and primary urothelial cells) cells from prostate (B&C), breast (D&E), and bladder (F&G) over multiple pulses (0, 1, 2, 5, & 10). H) Intrinsic resistance for indicated cell lines over the course of FSS exposure. Solid lines indicate transformed cells and dashed lines indicate non-transformed lines. Intrinsic resistance correlates with transformation status ($p<0.0001$; repeated measures three-way ANOVA). I) Membrane repair for the 3 paired cell lines over the course of FSS exposure for the same experiment shown in H. Membrane repair does not correlate with transformation status ($p>0.05$; repeated measures three-way ANOVA).

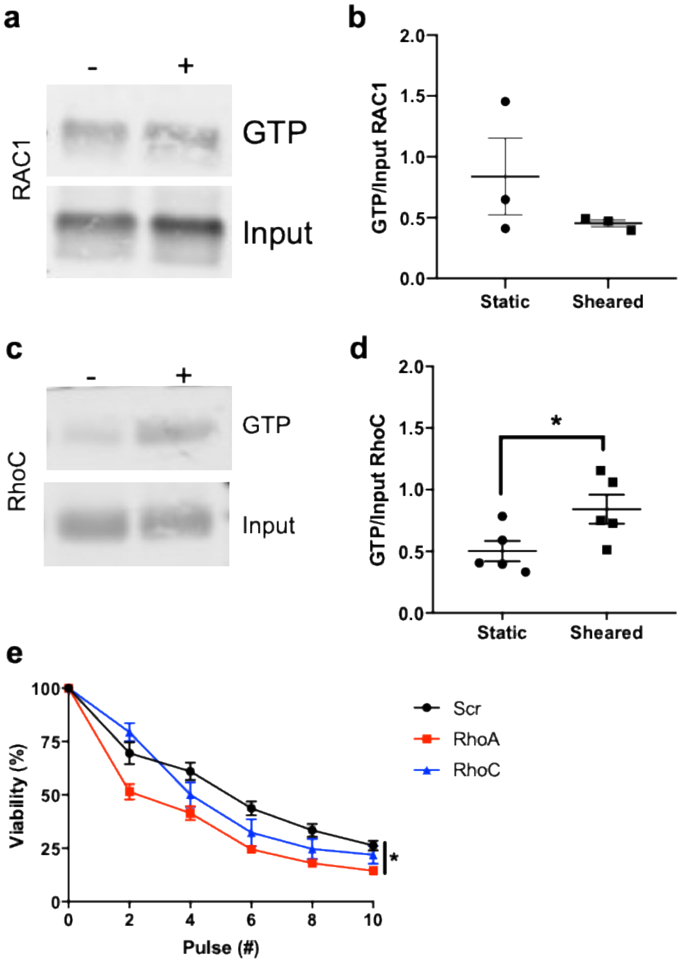


**Figure S3: Exposure to FSS activates RhoA/C and not RAC GTPases.** a. RAC1 pulldown blot for static (-) and sheared (+) cells and b. analysis of RAC1 activity assay indicating that GTP-RAC1 levels do not change in response to FSS ($p>0.05$, n=3, T-test). c. RhoC pulldown blot for static (-) and sheared (+) cells and d. RhoC activity assay data demonstrating an increase in GTP bound RhoC after FSS exposure ($p<0.05$, n=5, T-test). e. Viability after exposure to FSS for GS689.Li cells expressing shRNA against RhoA (n=6), RhoC(n=6), or non-targeting control (SCR) (n=12). RhoA knockdown cells have significantly reduced viability from pulse number 2 to 10 in comparison to SCR cells ($p<0.05$, t-test with Bonferroni correction), while RhoC does not cause a significant change in viability ($p>0.05$, t-test with Bonferroni correction).

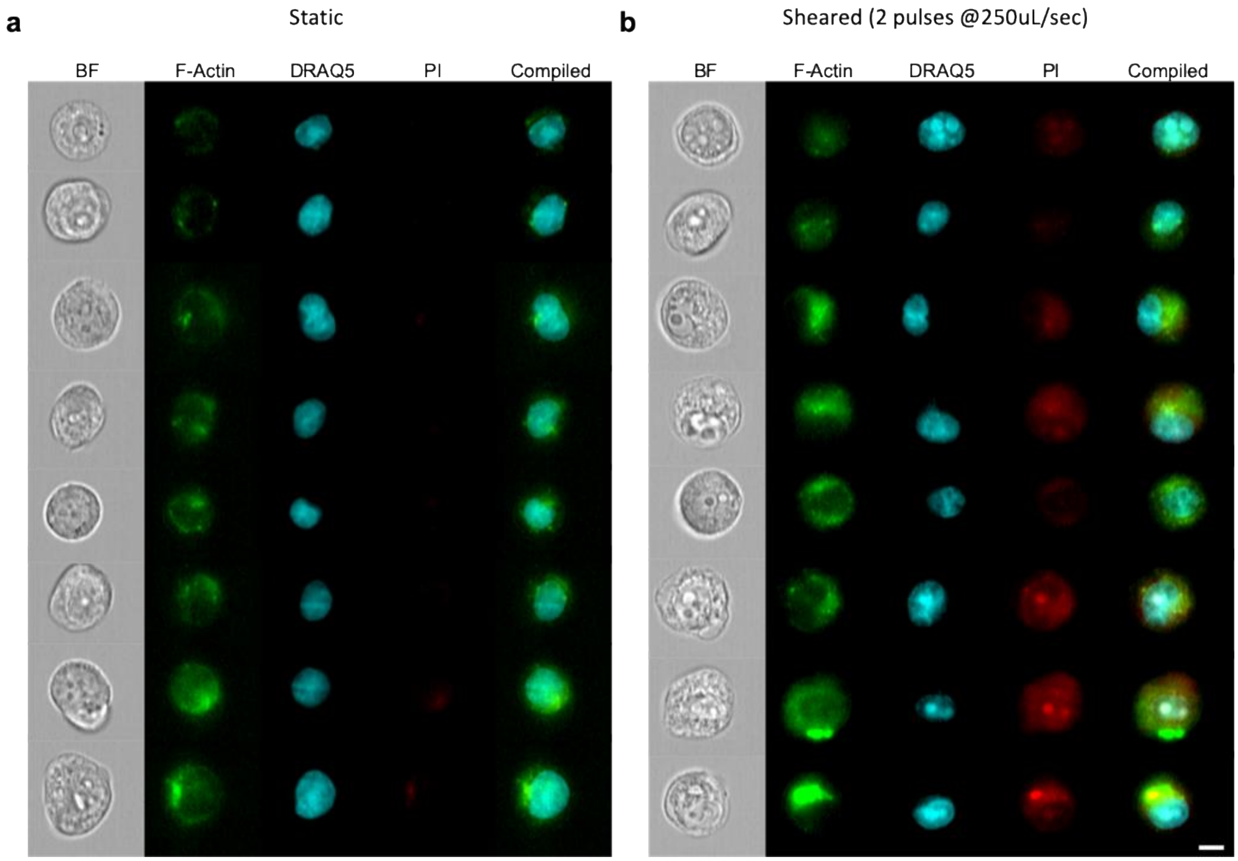


**Figure S4: Exposure to FSS results in increased cortical F-actin.** Representative images obtained by imaging flow cytometry A) Static control and B) cells exposed to 2 pulses of FSS, showing a clear increase in F-actin signal. Images are selected to represent the observe range of F-actin intensity: top two rows of images represent cells from the lower quartile; the middle 4 rows of images represent cells with median intensity of actin; and bottom two rows of images are representative of the upper quartile. Bar, 15 µm.

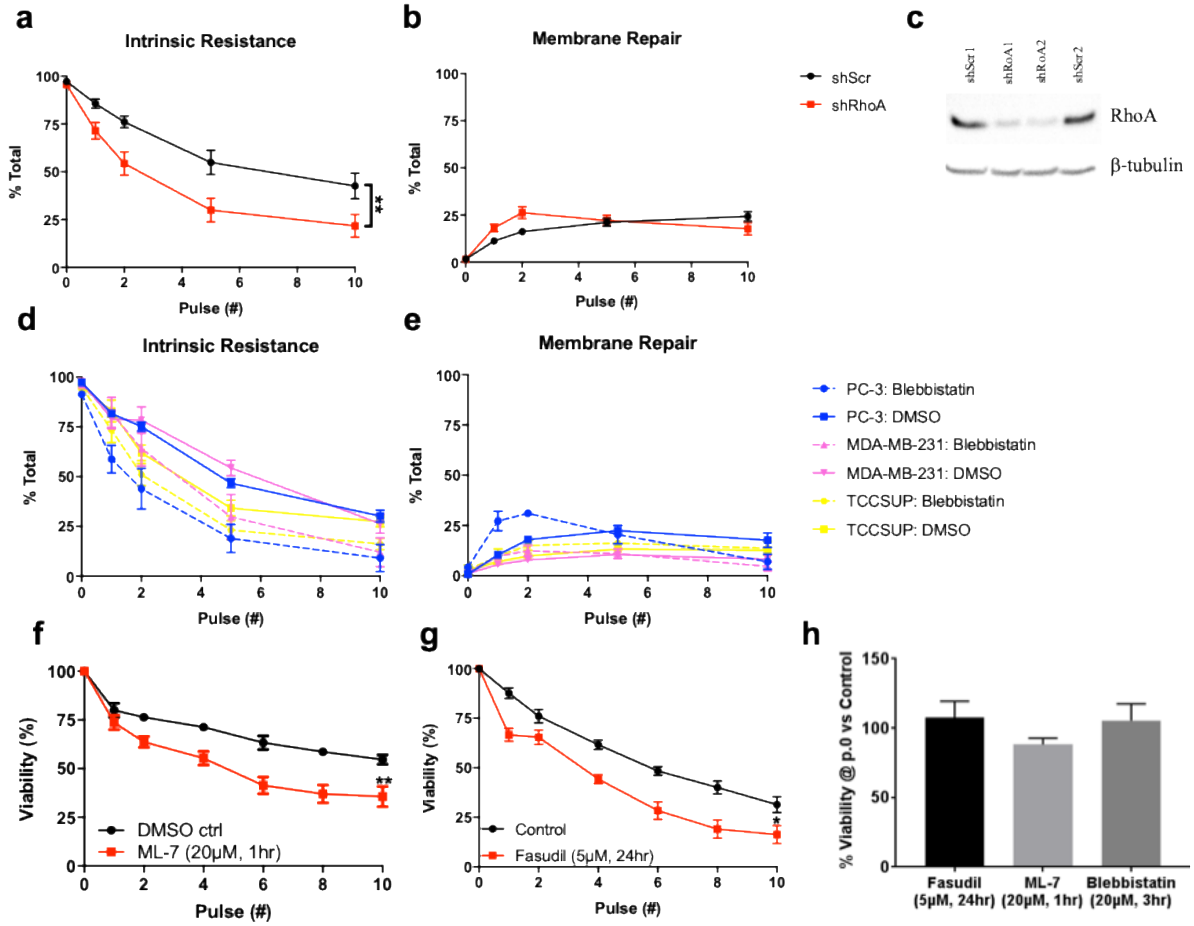


**Figure S5: RhoA-myosin II axis confers elevated intrinsic resistance to FSS.** A) Inhibition of RhoA significantly reduces intrinsic resistance compared to SCR control cells (**p<0.01; 2-way ANOVA), B) while the effects of RhoA knockdown on membrane repair is not statistically significant (p>0.05; 2-way ANOVA). C) Western blot of PC-3 cells with control shRNA and shRNA against RHOA, showing RhoA knockdown. D) A 3 way ANOVA of the intrinsic resistance data demonstrated that blebbistatin treatment accounted for a significant portion of the variability (p<0.01) and that the response to blebbistatin treatment did not depend on the cell line (p>0.05). Additionally, blebbistatin treatment influenced how the cells responded to each pulse of FSS (p<0.001). E) 3-way ANOVA of the membrane repair data demonstrates that membrane repair is different among cell lines (p<0.0001) and it is impacted by blebbistatin treatment (p<0.05). F) ML-7, myosin light-chain kinase inhibitor, (20µM, 1hr) sensitizes PC-3 cells to FSS (**p<0.01 2-way ANOVA). G) Pre-treatment with fasudil, a ROCK inhibitor, (5µM, 24 hr) sensitizes PC-3 cells to FSS (*p<0.05, 2-way ANOVA). H) None of these agents significantly reduced the viability of cells prior to FSS exposure.

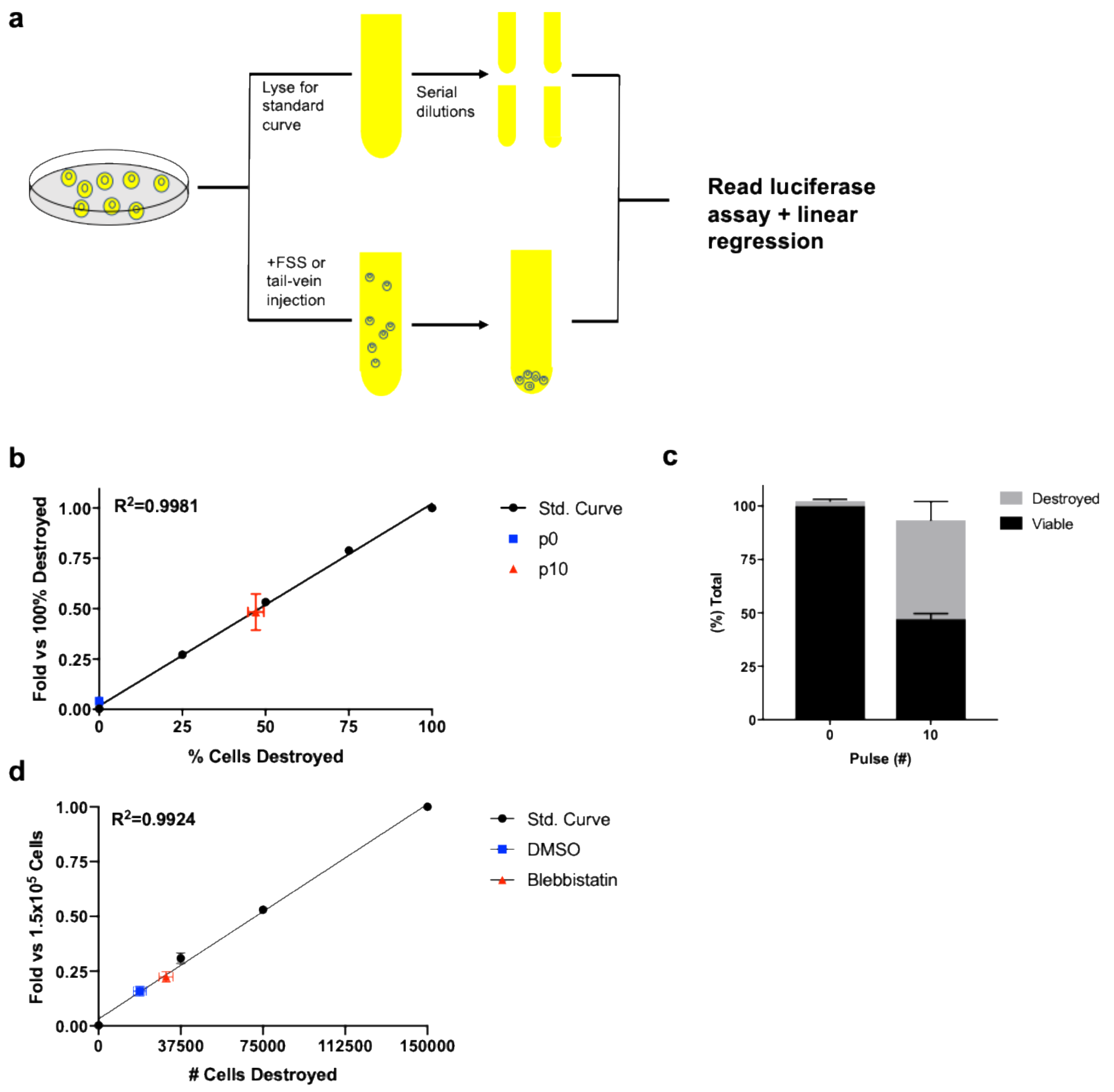


**Figure S6: Validation of assay for measuring cell destruction based on levels of cell-free luciferase.** A) Schematic of experimental workflow. B) Graph of the compiled data from the standard curves from lysed cells and supernatant from sheared cells used for validating the method with the in vitro FSS assay. C) Bar graph showing that the cell-free luciferase assay in (B) accurately accounts for the loss of viability from sheared cells ($p>0.05$; t-test). D) Standard curve and luciferase signal from the plasma of mice injected with cells compiled from 4 independent experiments, 2 with DMSO-treated cells expressing luciferase (n=7 mice) and 2 with blebbistatin-treated cells expressing luciferase (n=6 mice).

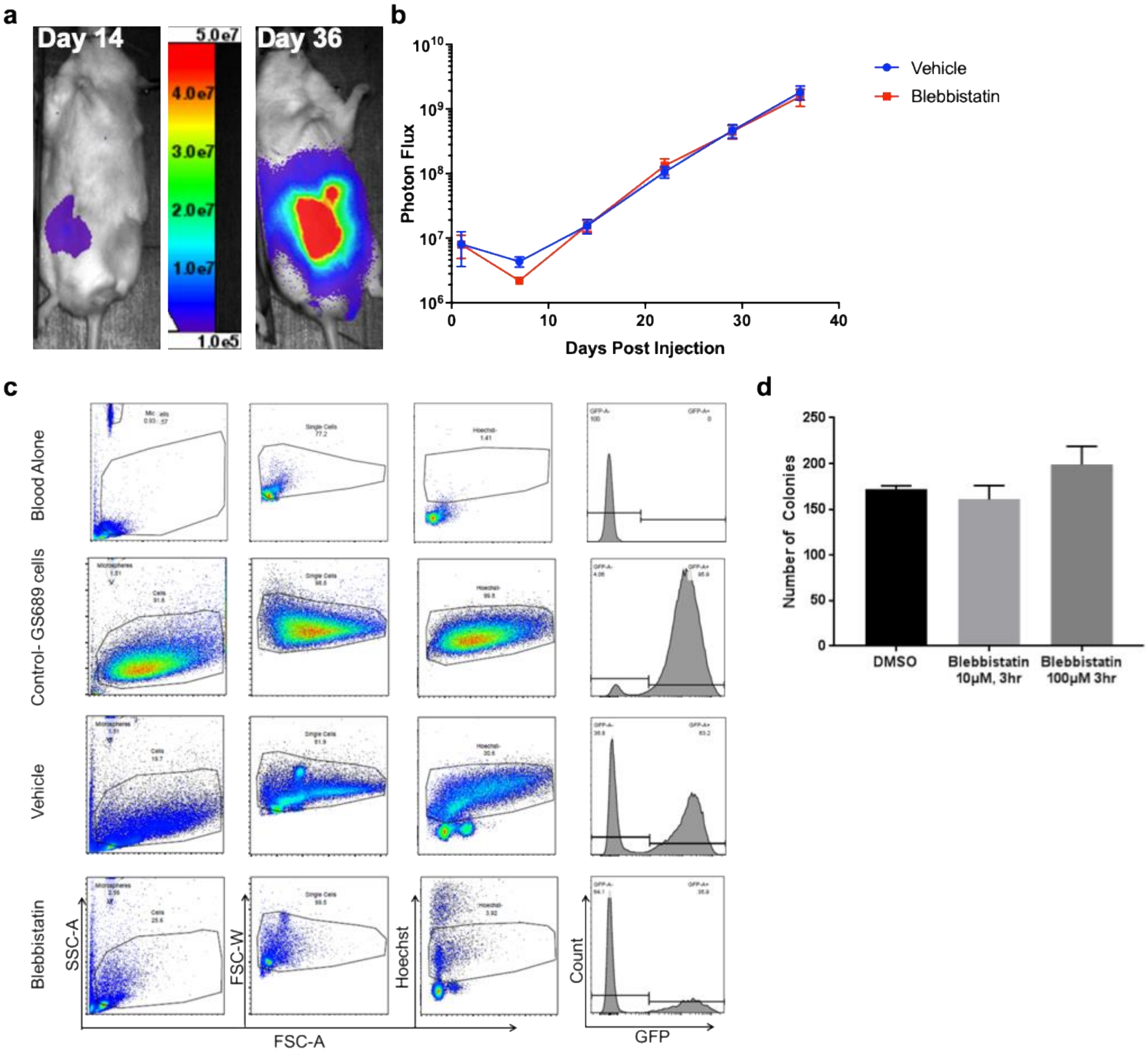


**Figure S7: Blebbistatin reduces steady-state CTC number**. A) Example BLI images and B) analysis of mouse tumor burden segregated by end group assignment, showing that tumor burden and growth did not different significantly between the vehicle and blebbistatin-treated groups ($p>0.05$; 2-way ANOVA). C) Flow analysis of blood alone, parental cells, blood from a vehicle treated mouse, and blood from a blebbistatin-treated mouse used to establish gating parameters. D) Clonogenic assay of GS689 cells treated with DMSO, 100μM (estimated maximum concentration) and 10μM of blebbistatin for 3 hours, demonstrating that the dose was not directly cytotoxic to cancer cells.

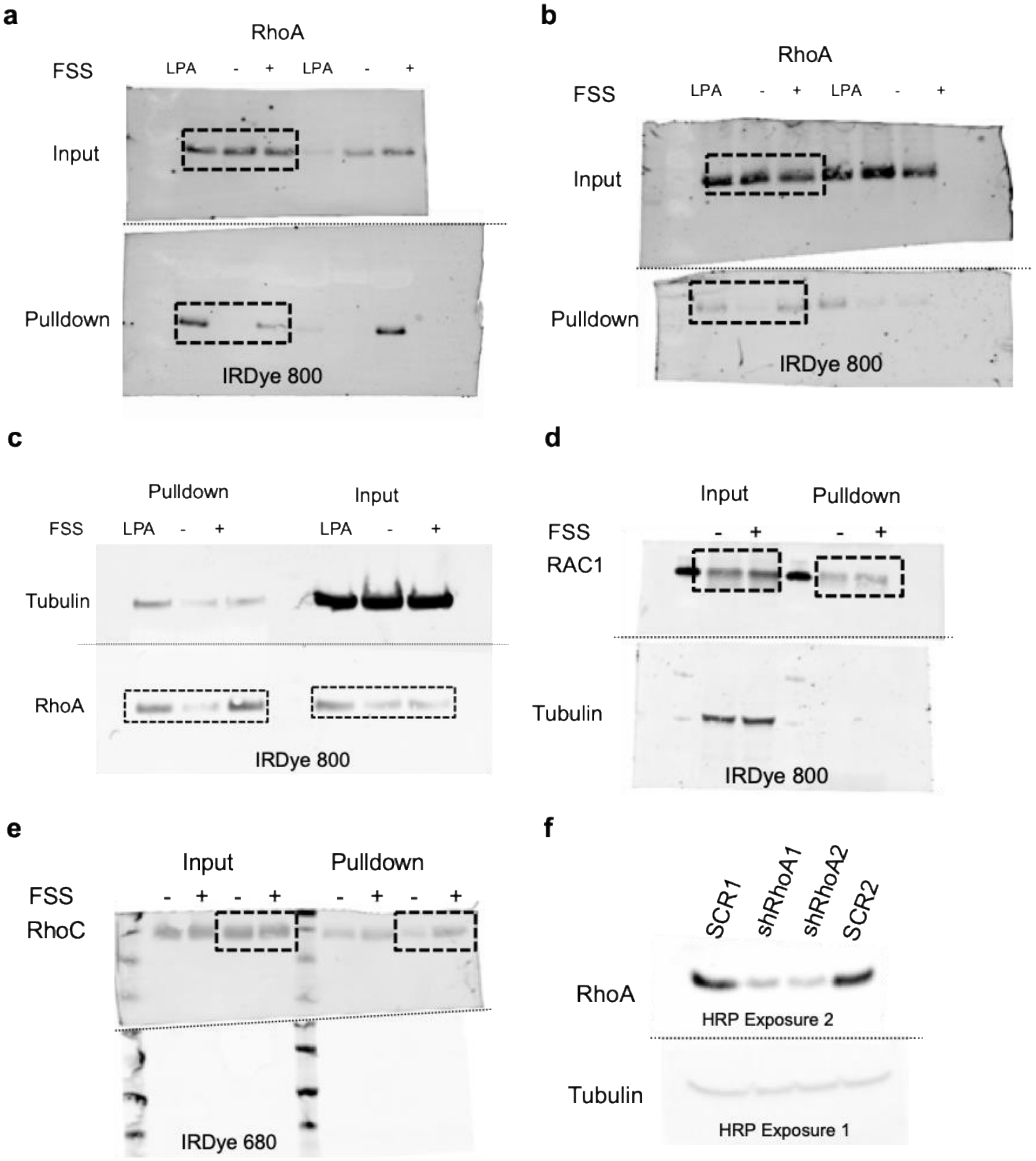


**Figure S8: Western blot scans presented throughout the paper.** Dotted line indicate where blot was cut to incubate with additional primary, dashed box indicate what is presented. Western for RhoA pulldown from a. PC-3 cells b. MDA-MB-231 cells, and c. TCCSUP cells. Western of PC-3 cells that have been exposed to FSS for d. RAC1 pulldown and e. RhoC pulldown. f. Western image of demonstrating RhoA knockdown. Exposure 1 was shorter than exposure 2

| Sample ID | DMSO | Blebbistatin | Microsphere Count | % Microsphere Inj. | % Inj. Cells Destroyed |
|---|---|---|---|---|---|
| **Control 1** | 677 | 658 | 46 | N/A | N/A |
| **Mouse 1** | 2993 | 2781 | 433 | 0.87% | 0.8% |
| **Mouse 2** | 3991 | 3359 | 518 | 1.04% | 2.6% |
| **Mouse 3** | 4577 | 3952 | 487 | 0.97% | 4.2% |
| **Mouse 4** | 2926 | 2461 | 328 | 0.66% | 4.1% |
| **Control 2** | 669 | 1103 | 61 | N/A | N/A |
| **Mouse 6** | 2311 | 2813 | 318 | 0.64% | 3.2% |
| **Mouse 7** | 3027 | 3743 | 427 | 0.85% | 6.1% |
| **Mouse 8** | 2473 | 3582 | 387 | 0.77% | 5.3% |

Table 1: **Fewer blebbistatin-treated cells lodge in the lung microvasculature.** Data from 2 independent animal experiments with colors corresponding to the dye used (Blue for Cell-Tracker Red and Green for Cell-Tracker Green), showing the count of nucleated cells as well as microspheres from whole lung section images. % injected cells destroyed column is based on cell-free plasma luciferase data and is colored to match the detection for which treatment group was measured.
